# Triceps Surae Muscle Ia Proprioceptive Weighting During Quiet Stance with Vision Occlusion

**DOI:** 10.1101/2025.06.25.661661

**Authors:** Gordon R. Chalmers

## Abstract

Visual, vestibular, proprioceptive and cutaneous sensory information is important for posture control during quiet stance. When the reliability of one source of sensory information used to detect self-motion for posture control is reduced, there may be a reweighting of inputs within and/or across the remaining sensory systems determining self-motion for postural control. Muscle vibration, which creates an illusion of muscle stretch and a compensatory movement to shorten the vibrated muscle, may be used to determine the weighting of muscle spindle Ia proprioception for posture control. The objective of this study was to determine the effect of vision occlusion on triceps surae muscle Ia proprioceptive weighting for postural control during quiet stance, utilizing 80 Hz muscle vibration and a quantitative measure of the body’s anterior to posterior ground center of pressure response to triceps surae muscle vibration in freely standing subjects. Subjects (N = 41; mean(standard deviation), 19.6(2.0) years) were examined as they stood with eyes open or eyes closed. Ground center of pressure was measured during quiet standing with, and without, bilateral vibration of the triceps surae muscles. The mean backward center of pressure shift induced by triceps surae vibration was significantly greater during the eyes closed condition compared to eyes open (eyes closed: -4.93(1.62) centimeters; eyes open: -3.21(1.33) centimeters; p = 6.85E-10; Cohen’s d = 1.29). Thirty-seven subjects increased, and two subjects decreased, their vibration induced center of pressure backward shift in the eyes closed condition compared to eyes open, although the magnitude of the change varied. Results support the idea that for most subjects, during an eyes closed stance there is an increased triceps surae muscle Ia proprioceptive weighting for postural control, due to the need for posture control to depend more on non-visual feedback.

## Introduction

Visual, vestibular, proprioceptive and cutaneous sensory information are important for the central control of steady upright standing (termed posture control in this study) during quiet stance [1–12]. For proprioceptive and cutaneous input, ankle sensory information plays an important role [1–13]. For ankle kinesthetic sense, muscle spindles are the major source of information, with skin receptors also playing a role [13–16]. When the reliability of one source of sensory information used to detect self-motion for posture control is reduced, there may be a reweighting (i.e. change in gain) of that and other sensory inputs utilized for posture control, within and/or across sensory modalities [10, 17, 18], although this does not always happen [19]. For example, when the reliability of ankle proprioception to predict body position during quiet stance is reduced by standing on a foam surface, causing the Ia signals from ankle muscles to no longer be a dependable indicator of body sway due to the confound of surface movements, there is a down weighting of effect (decrease gain) of the triceps surae (TS) muscle Ia signal, an up weighting of the effect (increase gain) of the lumbar paraspinal musculature Ia signal on posture control [20, 21] and an increased weighting of effect of the vestibular system on lower leg muscle activation [22]. Similarly, when surface sway referencing (moving the support platform in synchrony with the postural sway) reduces the ability for ankle proprioception to provide reliable information about body sway during quiet stance, there is an decreased gain of ankle proprioceptive input [23] with an increased gain of visual [7, 24], vestibulospinal [17, 23] and plantar somatosensory [25] input on posture control. Likewise, when the sensory information from the visual system is made increasingly less reliable for detecting body sway by progressively increasing the amplitude of visual surround sway referencing (moving everything in the visual field in synchrony with the postural sway) the effect of vision on posture control is progressively down weighted [17, 24, 26]. These examples demonstrate the reciprocal fashion with which the gain of some sensory channels are re-weighted when environmental conditions change, to ensure the maintenance of the appropriate muscle torques to sustain a stable stance [17, 27–29].

Muscle vibration may be used to determine the weighting of muscle proprioception for posture control during quiet standing [20, 21, 30, 31]. Muscle vibration causes an increase in the discharge rate of spindle primary (type Ia) endings and may also stimulate limited firing of spindle secondary endings, most Golgi tendon organs during muscle contraction, and adjacent cutaneous mechanoreceptors, with the dominant sensory response and effect via the Ia afferents [14, 32–34]. The increase in spindle firing in a vibrated muscle of a stationary limb creates an illusion that the muscle is being stretched [11, 34–37]. During quiet stance, this illusion in supporting muscles leads to a compensatory shift of the body’s center of mass (CoM) and ground center of pressure (CoP) in the direction to shorten the vibrated muscle. For example, when standing, TS muscle vibration results in an illusion of forward leaning with a resulting posterior shift of the CoP [20, 21, 30, 31, 38]. Similarly, tibialis anterior (TA) muscle vibration in a standing subject produces an anterior shift in the body’s CoP [31, 38, 39]. The shift in body position during vibration of supporting muscles is not due to autogenic excitation by the tonic vibration reflex [38] as there is a decrease in the electromyographic activity of the soleus muscle with vibration during stance [30, 38].

Differences in the magnitude of the vibration induced shift in a body’s CoP across testing conditions indicate differences in the weighting of the vibrated muscle’s Ia proprioceptive signal for standing posture control [20, 30, 31]. For example, the posterior CoP shift with TS muscle vibration during quiet stance is decreased when subjects stand on surfaces unstable in the sagittal plane, indicating that the TS muscle Ia proprioception influence on posture control has been down weighted, compared to stance on a solid surface [20, 21, 30, 39].

The visual system is an important source of afferent information for stabilizing sway during stance [1, 7, 17, 40]. The challenge of maintaining a steady posture is increased if a person is deprived of visual information, as demonstrated by an increased standing sway under eyes closed (EC) conditions [41–43]. Nevertheless, in healthy young adults, the removal of visual information for posture control usually does not lead to falling and has only small effects on posture stabilization [44, 45]. If visual information is not available for posture control have other sources of sensory information contributing to posture control, such as ankle Ia proprioception, been up weighted to compensate for the loss of vision? Proprioceptive signals from the leg muscles alone, without visual, vestibular or foot sensory input, can sufficiently maintain stable upright stance, as measured with an inverted pendulum [4].

Few studies have examined the weighting of the ankle muscle proprioceptive system for posture control during quiet stance when in an eyes open (EO) condition versus a condition lacking visual information. The effect of eye closure on leg proprioceptive weighting for postural control was examined by Peterka using continuously applied motion perturbation stimuli, finding an increased leg proprioceptive weighting during an EC compared to an EO condition during quiet stance [17]. The technique employed by Peterka is very effective at measuring overall leg proprioceptive weighting but is unable to address if the proprioceptive weighting of individual leg muscles is modified with eye closure. The muscle vibration technique may be used to examine the Ia proprioceptive weighing of individual muscles, as discussed above. Individual muscle examination is important because ankle musculature is considered to be a very important proprioceptive input for posture control [1, 2, 7].

It is unclear, however, if the muscle vibration technique demonstrates an increased ankle muscle Ia proprioceptive weighting during an EC condition, compared to EO, during quiet stance, as Peterka’s technique did for the whole leg [17], because the two existing relevant vibration studies have methodological limitations. Lackner and Levine [11] reported that during an EO condition in an illuminated room that a TS muscle vibratory illusion was not reported by subjects, while the vibratory illusion did occur in a dark room. These results suggest a lower weighting of the TS muscle Ia proprioceptive system on posture control during an EO condition, and a greater weighting when visual information is removed. Lackner and Levine [11], however, relied on qualitative reports by subjects of a body movement illusion occurring with vibration while the subjects were fixed to a back board, as opposed to quantitative measures of the body’s CoM or CoP response to muscle vibration in a freely standing subject. In contrast, Toosizadeh and coworkers examined the anterior to posterior (AP) CoM position in subjects, finding an shift in CoM during an EO but not an EC condition when gastrocnemius muscle vibration was applied [43]. This may demonstrate a greater weighting of gastrocnemius muscle Ia proprioception for posture control during an EO condition, compared to with EC. However, the 40 hertz (Hz) vibration frequency used by Toosizadeh and coworkers was selected for their study goal of examining the effects of low intensity vibration and was not optimal to examine Ia proprioceptive weighting, because spindle stimulation reduces as vibration frequency is reduced [3]. Forty Hz is only slightly above the approximately 30 Hz threshold for eliciting vibration induced postural responses in TA and soleus muscles in healthy adults [38], and is much lower than the approximately 80 Hz vibration frequency used in most Ia proprioception weighting studies discussed above, and which is the vibration frequency that is optimal to stimulate spindle primary endings [14, 46]. Further, the lack of a vibratory induced sway in the EC condition reported by Toosizadeh and coworkers may have been due to a rise in the threshold for discrimination of vibratory stimuli, which may occur with eye closure [43, 47], being revealed by their utilization of a stimulation frequence only slightly above the threshold, as opposed to a change in weighting of the Ia proprioceptive system. In summary, the findings of Lackner and Levine and those of Toosizadeh and coworkers appear contradictory, the former suggesting a higher TS muscle Ia proprioceptive weighting for posture control with an absence of visual information, and the latter suggesting a higher weighting during an EO condition. But the two studies used very different vibrations frequencies for stimulation and different outcome measures of subjective reports of a movement illusion in a subject fixed to a back board versus AP CoM shifts in a freely standing subject [11, 43], making direct comparison of results difficult.

The purpose of this study was to determine the effect of vision occlusion on TS muscle Ia proprioceptive weighting for postural control during quiet stance, utilizing 80 Hz muscle vibration and a quantitative measure of the body’s CoP response to muscle vibration in freely standing subjects. Eye closure eliminates the contribution of visual sensory cues to posture control [17]. It was theorized that while in the EC condition, compared to EO, the need for posture control to depend more on non-visual feedback [10, 17, 48, 49] would result in a greater weighting of TS muscle Ia proprioceptive information for posture control during quiet standing, and this would be measured by a greater shift in the body’s AP CoP being induced by TS muscle vibration in the EC than in the EO condition. Accordingly, the research hypothesis was that during quiet stance, the mean AP CoP shift in the direction of the bilaterally vibrated TS muscles would be greater with vision occlusion, than with EO.

## Materials and Methods

### Participants

A priori, a G*power calculation (version 3.1.9.7) determined that 34 subjects were needed to detect a moderate treatment effect (0.5) with an alpha error probability of 0.05 and a statistical power of 0.8 for the statistical analysis used (discussed below) [50] .

Subjects between the ages of 18 and 40 years of age with no neurological, vestibular, visual or lower limb musculoskeletal disorders, or neuromuscular medications were recruited, wearing of visual corrective lenses was allowed. The experiments were approved by the Western Washington University Institutional Review Board Human Subjects Ethics Committee according to the principles expressed in the Belmont Report and the Declaration of Helsinki, and subjects gave informed consent prior to testing by reading and then signing a written consent form. Recruitment and participation of subjects started on January 13, 2025 and ended on February 3, 2025.

### Experimental procedures and data acquisition

Subjects wearing shorts stood bare footed on a force plate (Balance Tracking Systems, San Diego, CA, U.S.A.) with arms loosely hanging at sides, feet shoulder width apart with a self-selected spay angle, while standing straight and relaxed without moving, in a quiet room with no distractions. Foot position was marked on the plate surface to minimize change if repositioning was required.

Vibrators, made using a DC electric motor with an offset mass (BestTong, China, model A00000531, https://www.besttong.co/) mounted within a solid tube (6 inches long, 1.5 inches diameter), powered through a motor controller (RioRand, China, model RR-MJLK-FBDG, https://www.riorand.com/) with a two-axis accelerometer (Dimension Engineering, Hudson, Ohio, United States, model DE-ACCM6G, https://www.dimensionengineering.com/products/de-accm6g) mounted on the tube, were attached bilaterally to the TS muscle at the base of the bulk of the gastrocnemius muscle to expose both the gastrocnemius and the soleus muscles to vibration [51]. Elastic straps were utilized to limit vibration of nearby tissues [52] (Fig 1). After attachment, each vibrator was calibrated to vibrate at 80 Hz by adjustment of the drive voltage of the motor while the frequency of the output of the accelerometer was monitored (digitized at 10,000 Hz, Micro 1401, Cambridge Electronic Designs, Milton, Cambridge England) and analyzed using Spike2 software (Cambridge Electronic Designs, Milton, Cambridge England). All vibration stimuli were simultaneously applied bilaterally. Standing subjects then received two seconds of vibration bilaterally to the TS muscles to experience the bilateral stimulation and not be startled when it started during trials. Prior to trials, to prepare subjects for possible unexpected sensations, they were instructed that the vibrators would vibrate their muscles and they may feel their body position shift, they were to let the body automatically handle its balance, not interfere with effects felt, not to resist any body tilt, and a researcher was positioned to prevent a fall if needed. Padded floor mats were placed around them for safety. Briefing subjects on what to expect during the experiment increases safety, and also increases the likelihood of them experiencing the vibration movement illusion [46, 53], but was not specific in direction to prevent intentional responses by subjects. Subjects were randomly assigned to the testing sequence of EO then EC, or EC then EO. For each trial, the subject adopted the EO or EC condition for 20 seconds before measures proceeded, to allow time for adaptation to the new sensory condition [54, 55] then force plate data collection was started. After 20 seconds vibration was then initiated and continued for 20 seconds (total 40 seconds of data collection).

**Fig 1.**
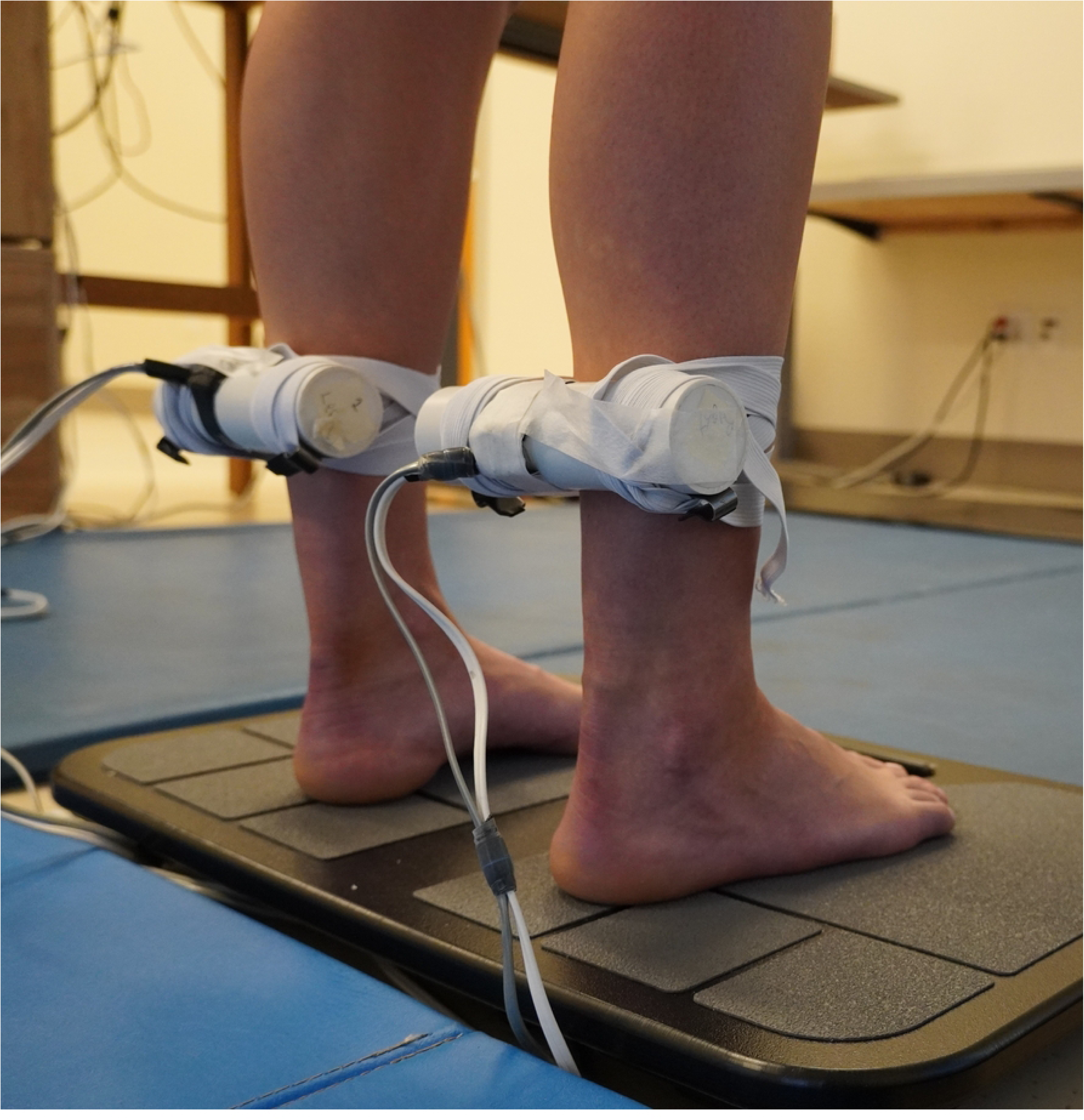
Placement of muscle vibrators. Vibrators were attached bilaterally to the TS muscle.

During the 40 second period BTrackS™ Assess Balance software (Balance Tracking Systems, San Diego, CA, U.S.A.) recorded the subject’s CoP on the force plate. Demura and co-workers examined standing sway frequency in healthy young people and demonstrated that the CoP power spectrum peak is at 0.2 Hz in both the AP and ML directions, there was 15% or less of the total power in the 2 - 10 Hz range, and that the mean power frequency was at 0.96 Hz and 1.06 Hz in the ML and AP directions, respectively [56, 57]. Similarly, Prieto and coworkers found that 95% of the power in a frequency analysis of standing sway occurs below 0.925 Hz with EO and 1.05 Hz with EC [58]. Accordingly, force plate data were collected at 25 Hz, as done in a prior similar study [9], and above the 3 Hz – 10Hz range which prior similar studies low pass filtered their data down to [20, 21, 31, 59–61]. During EO trials, subjects gazed straight ahead at a blank wall with a 95 centimeter (cm) by 60 cm colorful poster positioned at eye level at 300 cm distance. Subjects were instructed to select a spot in the poster to focus their gaze on. The use of a visual target was employed because this decreases standing sway during the EO test [62] and so enhanced the difference between the EO and EC conditions, compared to staring at a blank wall. The EC condition was produced by the subject closing their eyes, which was monitored. Diffused room lighting from above ensured that no source of light information could provide motion information during EC testing. In both the EO and EC conditions the head and eyes were held in the same position of head and eyes forward to minimize possible effects of head or eye movement on posture [63, 64]. If during a trial a subject moved more than the typical illusion generated body sway (e.g. subject moved an arm, foot, leg or torso), the trial was rejected and repeated up to once after a 5-minute seated rest period. To minimize a possible post-vibration carry over effect between trials [65], subjects had five minutes of seated rest between trials [43, 47], and each trial condition of EO and EC was conducted only once. The use of a single trial for each condition is common in muscle vibratory studies [20, 21, 43, 47, 59, 61, 66]. To minimize muscle thixotrophy effects, subjects stood from a seated rest position immediately before each trial, directly to their position on the force plate to produce a similar muscle contraction history across both conditions [67]. To ensure that the vibration frequency did not change once set, the above procedures were repeated six times in each of two subjects with EO during both trials. The frequency of the vibrators were recorded at set-up, during the first trial, and during the second trial, for each of the right (R) and left (L) legs. The CoP data from these trials for these two subjects was not recorded.

### Data analysis for each subject

CoP data were filtered using a 2nd order zero lag 4 Hz low pass Butterworth filter in the BTrackS™ Assess Balance software, then exported to an Excel file (version 2411, Microsoft Office 365, Redmond, WA U.S.A.). TS muscle vibration produces almost exclusively AP body movements [9], so only AP responses were analyzed for the CoP measures [3, 20, 21, 68]. The data collected were analyzed for a 15 second epoch in both the no vibration period (seconds 0-15) and the vibration period (seconds 25-40). The last 5 seconds of the no vibration period were omitted to ensure there was a time gap between the end of the no vibration analysis period and the start of the vibration. The first 5 seconds of the vibration period were omitted because in some subjects the vibration illusion can take several seconds to occur [11, 30, 39]. In each 15 second epoch analyzed the mean AP CoP was determined. Mean CoP reflects postural orientation [69]. For each of the two conditions of EO and EC, the AP CoP displacement induced by vibration, a reliable measure of vibration effects on postural sway [59], was calculated as: (*mean AP CoP during vibration – mean AP CoP during no vibration)*, positive value is forward and negative value is backward, so greater negative difference values indicate greater backward lean induced by vibration [20, 21, 31, 59]. The greater the magnitude of the vibration induced shift in the standing body’s CoP during muscle vibration, the greater the weighting of the vibrated muscle’s Ia proprioceptive signal for posture control [20, 30, 31].

### Statistical analyses

To determine if a subject responded to the vibration stimulus with a vibration induced shift in the AP CoP, Cohen’s d was calculated to compare the mean AP CoP during the EO no vibration and vibration conditions for the subject, with values meeting or exceeding, 0.2, 0.5, and 0.8, indicating small, medium, and large effects [70]. Only subjects who responded to the vibration stimulus during the EO condition with at least a small backwards vibration induced shift in their AP CoP were included in the subsequent data analysis. Data were tested for deviations from normality using the Shapiro-Wilk test. To determine the effect of eye closure on TS muscle Ia proprioceptive weighting, a repeated measures two-tailed t-test, and a Cohen’s d calculation, compared the mean AP CoP displacement induced by vibration during the EO and EC conditions within subjects, for all subjects combined. Differences in the vibration induced AP CoP shift across conditions tested indicate differences in the weighting of the vibrated muscle’s Ia proprioceptive signal for posture control [20, 30, 31]. For the tests of the vibrator frequency consistency across trials, the frequency data were analyzed using a two-way between (right and left legs) within (set-up, trial 1, trial 2) ANOVA. SPSS (version 29.0.2.0) and Microsoft Excel were used for calculations. A significance level was set at p < 0.05, and results are reported as mean(standard deviation).

## Results

Forty-one subjects were studied (age: 19.6(2.0) years; weight: 147(30) lbs; self reported as female = 25, male = 15, not female or male = 1). Two subjects did not respond to the vibration stimulus in the EO condition with at least a small backward shift of their AP CoP, based on Cohen’s d criterion. The remaining 39 subjects were included in the data analysis. Thirty-eight of these subjects had a large backward vibration induced AP CoP shift and one had a medium shift during the EO condition, based on Cohen’s d criteria. An example of a subject’s CoP record during the EO and EC trials, is shown in Fig 2. The data did not deviate from normality. The mean backward CoP shift induced by TS vibration was significantly greater during the EC condition compared to EO, the difference being large, based on Cohen’s d criterion (EC: -4.93(1.62) cm; EO: -3.21(1.33) cm; p = 6.85E-10; Cohen’s d = 1.29) (Fig 3). Thirty seven of the 39 subjects increased their vibration induced AP CoP backward shift in the EC condition, compared to EO, although the magnitude of the increase varied, reflected by the range of downward slopes of the thin solid lines in Fig 3. In contrast, two subjects decreased their vibration induced AP CoP backward shift in the EC condition, compared to EO, shown by two upward sloped thin solid lines in Fig 3. One subject who did not respond to the vibration during the EO condition with a backward shift of the AP CoP did demonstrate a large backward shift during the EC condition, based on Cohen’s d criteria (thin downward sloped dashed line, Fig 3), indicating this subject also increased their TS muscle Ia proprioceptive weighting with EC, although they could not be included in the calculation of the group means because they did not respond to the vibration during the EO test.

**Fig 2.**
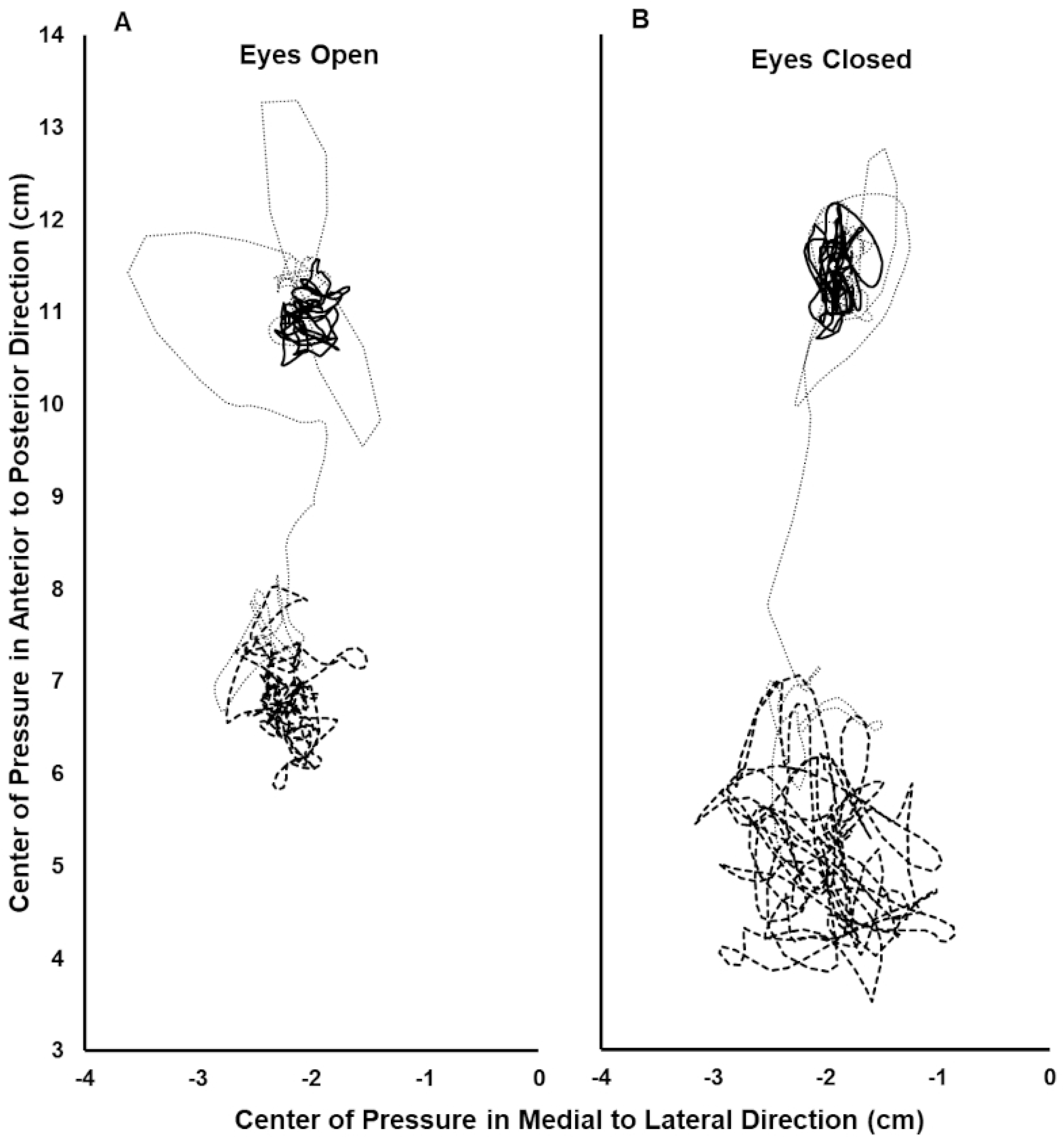
Center of pressure (CoP) record during the eyes open (A) and eyes closed (B) trials for a single subject. The CoP path during the initial 15 second period analyzed with no vibration (solid line), the final 15 second period analyzed with vibration (thick dashed line), and the 10 second interval between these periods which was not included in the analysis (thin dotted line) is shown. In the anterior to posterior direction, greater positive values are further anterior. Note the greater posterior shift with vibration in the eyes closed (B), versus the eyes open (A), condition.

**Fig 3.**
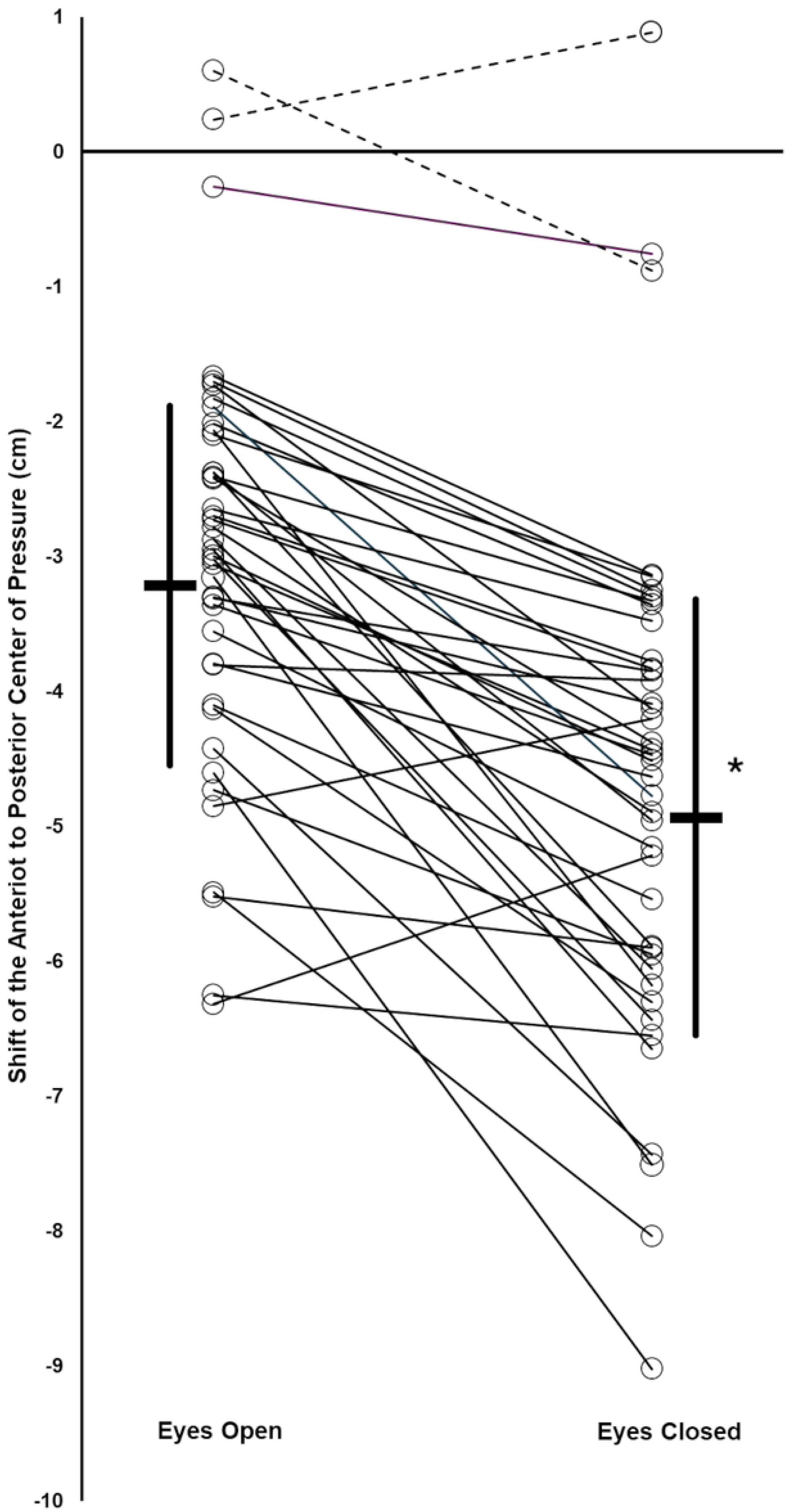
Shift in the anterior to posterior (AP) center of pressure (CoP) of subjects with triceps surae muscle vibration, during eyes open (EO) and eyes closed (EC) conditions. Positive value is forward and negative value is backward, so greater negative values indicate greater backward lean induced by vibration. Thick horizontal bars indicate group mean values, with standard deviation (thick vertical lines) for EO and EC conditions, for subjects who responded to the muscle vibration during the EO condition with at least a small backward CoP shift. For each subject, their AP CoP shift during EO and EC conditions are shown by open circles, with lines connecting the EO and EC data for the same subject, thin solid lines for subjects who responded to the muscle vibration during the EO condition with at least a small backward CoP shift, thin dotted lines for the two subjects who did not responded to the muscle vibration during the EO condition with at least a small backward CoP shift. * significant mean difference between EO and EC conditions: p = 6.85E-10; Cohen’s d = 1.29.

For the tests of vibrator frequency consistency across the trials, examined in two subjects, there was no significant change in vibration frequency over the course of the experiment, or when comparing the two legs (Set-up: R = 79.5(0.4) Hz, L = 79.5(0.4) Hz; trial 1: R = 79.4(0.7) Hz, L = 79.6(0.5) Hz; trial 2: R = 79.4(0.5) Hz, L = 79.6(0.5) Hz, interaction p = 0.45, trial main effect p = 0.96, leg main effect p = 0.36).

## Discussion

The present study was conducted to determine if vision occlusion increases TS muscle Ia proprioceptive weighting for postural control during quiet stance. The novel contribution of this study was that it was the first to examine this question utilizing an 80 Hz muscle vibration, optimal for stimulation of muscle spindles, and a quantitative measure of the body’s CoP response to muscle vibration in freely standing subjects. Prior studies examining other testing conditions and sensory systems demonstrate that when one source of sensory information used to detect self-motion for posture control is reduced, there may be an increased weighting of the effect of other sensory inputs contributing to posture control, within and/or across sensory modalities [10, 17, 18]. The present data demonstrate a large (Cohen’s d definition) increase in mean TS muscle Ia proprioceptive weighting for postural control during quiet stance when eyes are closed, compared to the EO condition (Fig 3).

Two previous studies reported an increase in proprioceptive weighting from the leg during EC versus EO conditions [11, 17]. Direct comparison of the magnitude of the increased TS muscle Ia proprioceptive weighting under EC conditions between these two prior studies and the current study is not possible, however, because Peterka utilized a very different procedure employing continuously applied motion perturbation and determined whole leg proprioception, while Lackner and Levine relied on qualitative reports by subjects of a body movement illusion occurring with TS muscle vibration [11, 17]. Toosizadeh found a vibration induced shift in CoM during an EO, but not an EC condition, possibly indicating a greater weighting of gastrocnemius muscle Ia proprioception for posture control during EO than during EC, opposite to the results of the present study [43]. Toosizadeh and coworkers, however, utilized a 40Hz muscle vibration frequency which was only slightly above the threshold for eliciting vibration induced postural responses in soleus and TA muscles [38] and so may not be optimal for examining the research question of the present study. Additionally, the lack of a vibration effect on posture during the EC condition which they observed may have been due to a rise in threshold for discrimination of vibratory stimuli occurring with eye closure [43, 47] , as opposed to a lower Ia proprioceptive weighting with eye closure compared to during the EO condition.

Hypothesizing an eye closure induced change in ankle proprioceptive weighting for posture control is reasonable due to the importance of proprioception, particularly from the ankle, for posture control [7, 9, 13]. Selection of the ankle plantar flexor muscles as the targets for measurement of ankle proprioceptive feedback during eye closure is appropriate because standing sway increases under EC conditions [41, 42], with AP sway, the predominate direction of effect of the TS muscles being tested, increasing more than medial lateral sway [40–42, 71, 72]. Future studies examining TA muscle Ia proprioceptive weighting with eye closure would be interesting, given the possibility that the TA muscles may be even more important than the TS muscles for providing AP proprioceptive feedback from the ankle [5, 13]. The mechanism by which TS muscle Ia proprioceptive weighting was changed in subjects during EC stance was not examined. It is likely that it is not due to a change in muscle spindle sensitivity in the TS muscles, because eye closure during stance does not change the fusimotor control of pre-tibial muscles [73]. Therefore, an alteration in Ia proprioceptive information central processing likely occurs [31].

The response to muscle vibration varied widely between subjects. The amount of backward lean induced by TS muscle vibration with EO was expected to vary between subjects, because intersubject variability in the magnitude of the muscle vibration illusion effect has been previously reported, including subjects who fail to respond with any vibration induced body movement [11, 30, 35]. A novel finding in this report is the wide variability in the increase in TS muscle Ia proprioceptive weighting with eye closure, compared to EO, for most subjects and the observation that two subjects decreased TS muscle Ia proprioceptive weighting with eye closure, reflected by the range of slopes of the thin solid lines in Fig 3. Isableu and coworkers have previously established that individuals vary in their degree of dependence on the visual field for body posture control [49].

Individuals classified as highly visual field independent subjects were better able to use nonvisual information from proprioceptive and vestibular sources for posture control, while highly visual field dependent subjects had difficulties utilizing these nonvisual cues for posture control, when challenged with reduced visual cues of self-orientation and self-motion [49]. Accordingly, intersubject differences in the degree of dependence on the visual field for body posture control, and the associated range in abilities to use nonvisual information from proprioceptive and vestibular sources for posture control, may explain the varying degrees and directions of changes in the TS muscle Ia proprioceptive weighting when comparing EO and EC conditions.

### Limitations

The vibration applied was intended to stimulate ankle TS muscle spindle Ia proprioceptive feedback. But leg cutaneous stimulation also affects proprioception from the ankle joint [15, 16] and this stimulation would be influenced by the straps holding the muscle vibrators and the vibration employed. But leg cutaneous stimulation would have been the same during both the EO and the EC trials, and so would not be the cause of the difference in mean AP CoP backward shift with vibration when comparing the EC and EO conditions. The current study utilized the test measurement of vibration induced backward body sway. But backward body sway changes plantar foot pressure distribution, and plantar somatosensory feedback also contributes to posture control [15, 74]. Accordingly, there may have been an influence of a change in plantar somatosensory feedback during the vibration testing procedure which then influenced the degree of backward sway measured during both the EO and EC conditions. Eye movement during EO conditions affects posture control [63, 64], but this was minimized or eliminated by having the subjects focus on one visual target during EO testing, and observations of the subjects. It is unlikely that any eye movement during EC testing would affect the results, because saccades when the eyes are closed do not affect postural sway [75]. The current study utilized eye closure as the method to modify visual feedback to posture control. Eye closure is equally effective as the utilization of sway referencing visual surround conditions for eliminating the contribution of visual feedback to posture control [17]. Eye closure is not, however, the equivalent of utilizing a dark room with EO to eliminate visual feedback to posture control. In a dark room with eyelids open there are smaller displacements for a subject’s vertical projection of the center of gravity and for the difference between the CoP and the vertical projection of the center of gravity, compared to when standing with eyelids closed [72]. The present results may, therefore, not apply to dark conditions with open eyes.

## Conclusions

For most subjects, and as a group mean change, there was an increased TS muscle Ia proprioceptive weighting for postural control during an EC quiet stance, compared to an EO stance. Results support the idea that during an EC stance most individuals are able to depend more on TS muscle Ia proprioception for posture control, to partially compensate for the lack of visual information.

## Acknowledgements

Dr. Harsh Buddhadev is thanked for comments on the manuscript.

